# stPlus: a reference-based method for the accurate enhancement of spatial transcriptomics

**DOI:** 10.1101/2021.04.16.440115

**Authors:** Shengquan Chen, Boheng Zhang, Xiaoyang Chen, Xuegong Zhang, Rui Jiang

## Abstract

**Motivation:** Single-cell RNA sequencing (scRNA-seq) techniques have revolutionized the investigation of transcriptomic landscape in individual cells. Recent advancements in spatial transcriptomic technologies further enable gene expression profiling and spatial organization mapping of cells simultaneously. Among the technologies, imaging-based methods can offer higher spatial resolutions, while they are limited by either the small number of genes imaged or the low gene detection sensitivity. Although several methods have been proposed for enhancing spatially resolved transcriptomics, inadequate accuracy of gene expression prediction and insufficient ability of cell-population identification still impede the applications of these methods.

**Results:** We propose stPlus, a reference-based method that leverages information in scRNA-seq data to enhance spatial transcriptomics. Based on an auto-encoder with a carefully tailored loss function, stPlus performs joint embedding and predicts spatial gene expression via a weighted *k*-NN. stPlus outperforms baseline methods with higher gene-wise and cell-wise Spearman correlation coefficients. We also introduce a clustering-based approach to assess the enhancement performance systematically. Using the data enhanced by stPlus, cell populations can be better identified than using the measured data. The predicted expression of genes unique to scRNA-seq data can also well characterize spatial cell heterogeneity. Besides, stPlus is robust and scalable to datasets of diverse gene detection sensitivity levels, sample sizes, and number of spatially measured genes. We anticipate stPlus will facilitate the analysis of spatial transcriptomics.

**Availability:** stPlus with detailed documents is freely accessible at http://health.tsinghua.edu.cn/software/stPlus/ and the source code is openly available on https://github.com/xy-chen16/stPlus.

**Supplementary information:** Supplementary data are available at *Bioinformatics* online.

## 1 Introduction

Recent advances in single-cell RNA sequencing (scRNA-seq) techniques have stimulated efforts to unravel the heterogeneity of cell types, revolutionized the understanding of various complex tissues, and boosted the development of modern cellular and molecular biology. However, scRNA-seq requires a dissociation step to obtain cell suspension, leading to a loss of spatial context. The maintenance of spatial context is crucial for understanding cellular characteristics and reconstructing tissue architecture in normal physiology or under perturbation.

To elucidate single-cell heterogeneity and define cell types while also retaining spatial information, a number of remarkable methodologies have been recently developed to profile spatially resolved transcriptomics. These methodologies can be grouped into two main categories: 1) se-quencing-based technologies that provide unbiased capture of the transcriptomic landscape via capturing and quantifying the mRNA population of molecules in situ, such as ST (Stahl, et al., 2016), HDST (Vickovic, et al., 2019) and Slide-seq (Rodriques, et al., 2019); and 2) imaging-based technologies that offer higher spatial resolutions by fluorescence in situ hybridization (FISH) or in situ sequencing, such as osmFISH (Codeluppi, et al., 2018), seqFISH+ (Eng, et al., 2019), MERFISH (Moffitt, et al., 2018), and STARmap (Wang, et al., 2018). These approaches are often complementary and differ in their target throughput, coverage, and spatial resolution. For example, sequencing-based methods offer relatively high throughput but have been limited by spatial resolution, while imaging-based methods can provide subcellular resolution but have been limited in terms of sequence coverage and overall throughput (Larsson, et al., 2021; Zhuang, 2021). In this study, we focus on the imaging-based methods with the understanding that increasing spatial resolution can help define transcriptomic gradients within tissues more accurately, and allow detection of the subcellular localization of transcripts (Chen, et al., 2021).

In imaging-based methodologies, early methods based on FISH, such as osmFISH, can provide excellent signal quality in tissue imaging with even higher gene detection sensitivity than scRNA-seq (Codeluppi, et al., 2018). Nevertheless, the molecular crowding problem presents a challenge to scaling up the number of genes imaged. Although recent advanced technologies can help mitigate the molecular crowding problem and image more than 10,000 genes in individual cells (Eng, et al., 2019; Xia, et al., 2019), increasing the number of genes imaged will, in general, lead to increment of the overall measurement time and/or reduction of the measurement accuracy for multiplexed FISH and in situ sequencing (Zhuang, 2021). Therefore, a more general challenge for imaging-based profiling of spatially resolved transcriptomics is to balance the number of genes imaged and the gene detection sensitivity. As suggested by Wang, et al., a conceptually simple approach based on the idea of divide-and-conquer is to divide the targeted genes into multiple groups and image them sequentially one group at a time (Wang, et al., 2018). Moreover, computational approaches can help fill this gap by incorporating scRNA-seq data as reference and predicting genome-wide expression of cells profiled with a targeted gene set (Zhuang, 2021), given that recent efforts of cell atlas consortiums have generated massive amounts of scRNA-seq data.

Computational methods for enhancing spatially resolved transcriptomics are anticipated to accurately predict expression of unmeasured genes on the data with small number of genes imaged, and effectively impute expression of imaged genes to better identify cell populations on the data with low gene detection sensitivity. Existing computational methods can be divided into two major categories: 1) probabilistic methods, such as gimVI (Lopez, et al., 2019), that generatively model spatial transcriptomic data while integrating reference scRNA-seq data by domain adaptation; and 2) joint embedding-based methods, such as Seurat (Stuart, et al., 2019) and Liger (Welch, et al., 2019), that perform joint dimensionality reduction for spatial data and reference data, and then impute the unmeasured genes in spatial transcriptomic data based on the linkage between cells in these two datasets. Most recently, SpaGE performs linearly joint embedding using genes shared between spatial and scRNA-seq datasets, and then predicts spatial gene expression by a *k*-nearest-neighbor (*k*-NN) approach (Abdelaal, et al., 2020). For the joint embedding-based methods, the first step plays the most important role in the enhancement of spatial transcriptomic data (Abdelaal, et al., 2020), while the second step is usually based on a general idea that integrates the information of neighboring cells in reference scRNA-seq data (Abdelaal, et al., 2020; Welch, et al., 2019). Abdelaal, et al. also provided an evaluation of the existing methods for the first time, and demonstrated that SpaGE achieves overall state-of-the-art performance. However, the use of only a certain fraction of the features (i.e., genes present in both datasets) cannot take full advantage of the reference scRNA-seq data, and thus limits the performance of joint embedding and final enhancement. Besides, although evaluation based on Spearman correlation coefficients can assess the prediction accuracy to some extent, it may not be the optimal evaluation metric because the Spearman correlation coefficients are relatively low in most instances even the visual inspection shows good enhancement for genes with known spatial pattern (Abdelaal, et al., 2020; Lopez, et al., 2019). In addition, Spearman correlation coefficients cannot reflect performance for the identification of cell populations using enhanced spatial transcriptomic data.

Motivated by the above understanding, we propose stPlus, a reference-based method for the accurate enhancement of spatial transcriptomics. stPlus is built upon an auto-encoder with a carefully tailored loss function for leveraging the holistic information in reference scRNA-seq data instead of only genes shared with spatial transcriptomic data. With the learned cell embeddings, stPlus predicts gene expression in spatial transcriptomics via a weighted *k*-NN approach. We also introduce a cluster-ing-based approach to assess the cell heterogeneity maintained in the predicted spatial profiles by metrics suitable for different scenarios. We conduct a comprehensive evaluation by gene-wise and cell-wise Spearman correlation coefficients and four metrics for cell clustering. Benchmarked against state-of-the-art methods across a variety of spatial and scRNA-seq dataset pairs, stPlus provides superior enhancement performance and can scale to large datasets. We also demonstrate that the predicted spatial gene expression can offer comparable or even better performance to identify cell populations than the measured spatially resolved transcriptomics. We anticipate stPlus will help mitigate the technical limitations and better characterize the transcriptomic pattern of complex tissues.

## 2 Materials and methods

### 2.1 The model of stPlus

stPlus aims at enhancing spatial transcriptomics by accurately predicting expression of unmeasured genes and effectively imputing expression of measured genes. The input of stPlus is the target spatial data and the reference scRNA-seq data profiled from matching or similar tissue as the spatial data. These two data can be represented by two gene-by-cell matrixes, respectively. Note that the cells between these two data are not matched, and the genes in reference data usually include most of the genes in spatial data. Users can specify any genes from the reference data to be predicted. The output of stPlus is a gene-by-cell matrix containing the predicted expression of each specified gene for each cell in the spatial data.

The enhancement process of stPlus can be divided into three main steps: 1) data processing to prepare for joint embedding; 2) joint embedding of the single cells in spatial transcriptomic data and reference scRNA-seq data; and 3) predicting the expression of spatially unmeasured genes based on the cell embedding and reference scRNA-seq data (Fig. 1).

**Fig. 1.**
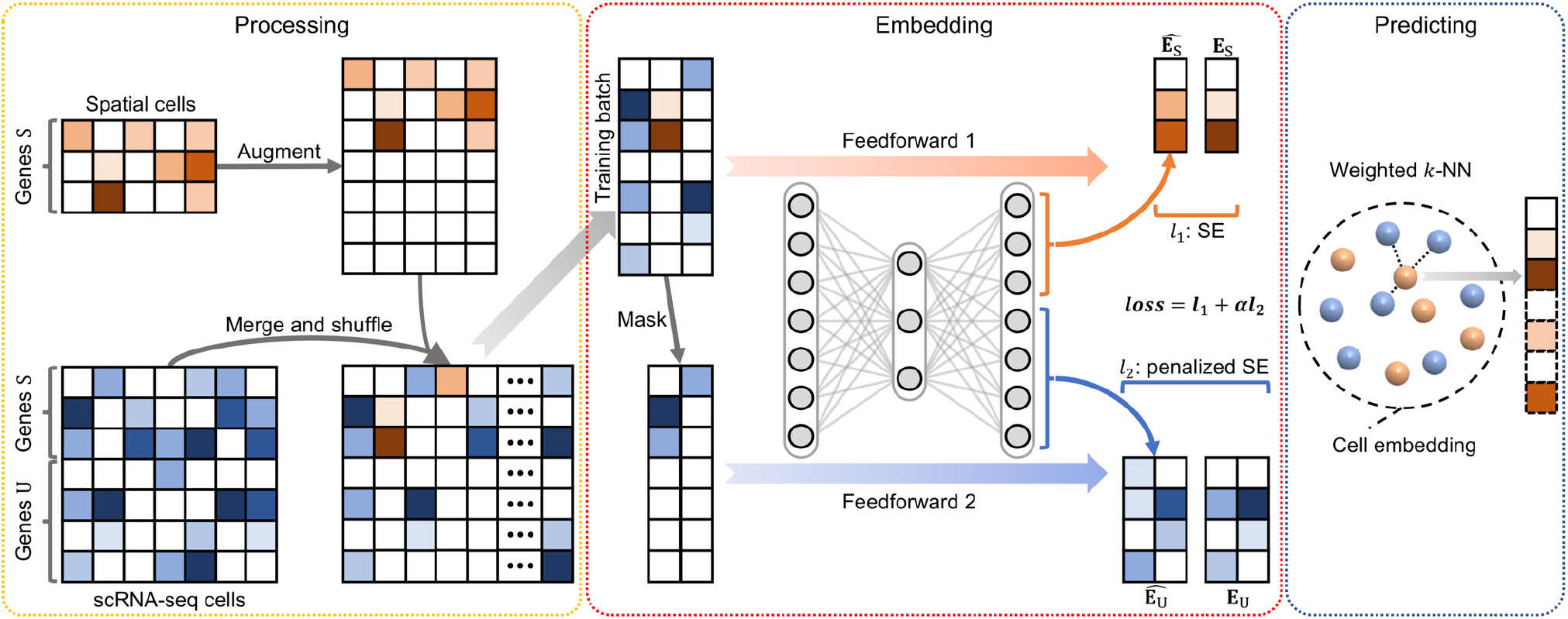
A graphical illustration of the stPlus model. stPlus first augments spatial transcriptomic data and combines it with reference scRNA-seq data. The data is then jointly embedded using an auto-encoder. Finally, stPlus predicts the expression of spatially unmeasured genes based on weighted *k*-NN.

In the data processing step, to avoid introducing highly noisy genes, stPlus selects the top 2,000 highly variable genes from genes only present in scRNA-seq data, as suggested by the widely-used toolkits for scRNA-seq analysis (Stuart, et al., 2019; Wolf, et al., 2018). The set of selected genes is denoted as U, while the set of genes shared between spatial transcriptomic data and reference scRNA-seq data is denoted as S. For the spatial transcriptomic data, stPlus augments expression measurements of genes U with zeros, and unifies the gene order with that of the processed scRNA-seq data. Then, the spatial transcriptomic and scRNA-seq datasets are merged together, shuffled across cells, and fed into the next step.

The joint embedding step aims to fit each spatial transcriptomic cell to the most similar scRNA-seq cells, but not to perform single-cell data integration and batch-effect correction (Tran, et al., 2020), or dimensionality reduction for downstream analyses such as data visualization and cell clustering (Sun, et al., 2019). The joint embedding step plays the most crucial role in the enhancement of spatial transcriptomic data (Abdelaal, et al., 2020), while the state-of-the-art method performs linearly embedding without incorporating the information of genes unique to scRNA-seq data, which constitutes major hindrances to the enhancement performance. In this step, stPlus uses an auto-encoder with a carefully tailored loss function to incorporate the biological variation of scRNA-seq-unique genes.

More specifically, the auto-encoder projects the gene expression data with a dimension of S + U into a 1,000-dimension latent space with a fully-connected layer. The latent embedding is then decoded to the original space with another fully-connected layer. Both the layers use the rectified linear (ReLU) activation function, which outputs zero for negative values, to achieve non-linear transformation and overcome the vanishing gradient problem. As the essence of stPlus, the loss function consists of two parts: 1) loss of the reconstruction for shared genes S in the spatial transcriptomic data; and 2) data sparsity-penalized loss of the prediction for genes U in the reference scRNA-seq data.

To access the two parts of loss function, stPlus feedforwards each training batch twice, respectively. Firstly, stPlus feeds the batch into the auto-encoder, and extracts the decoded results corresponding to genes S for the spatial transcriptomic cells (denoted as 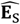)With the original expression data of genes S in the spatial transcriptomic cells (denoted as **E**_S_), the first part of loss function is calculated as follows:

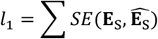

where *SE*(·,·) denotes squared error (squared L2 norm) between each element in the two inputs. Secondly, to mimic the augmented spatial transcriptomic data, stPlus extracts scRNA-seq cells in the batch, and masks the entries of genes U by setting the expression of genes U (denoted as **E**_U_) to zero. The masked scRNA-seq data is then fed into the auto-encoder, and the decoder output corresponding to genes U (denoted as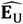) is re-garded as the prediction result for genes U in scRNA-seq cells. The ele-ment-wise squared error can thus be obtained by 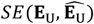. Given that scRNA-seq data usually contains technical artifacts (e.g., dropouts) and does not reflect the actual expression level, stPlus further penalizes 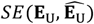 by down-weighting the errors of zero elements in **E**_U_ with the percentage of non-zero elements in reference scRNA-seq data (denoted as Q), i.e., one minus data sparsity. This strategy is based on the idea that the sparser the scRNA-seq data is, the more likely the zero elements are false negatives, and the prediction error should be penalized with lower weights. The second part of loss function is then computed as the following formula:

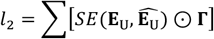

where ⊙ denotes element-wise multiplication, **Γ** denotes a matrix with the same dimension as **E**_U_, in which an entry is set to Q if the corresponding entry in **E**_U_ is zero, and is set to 1 otherwise. The two parts of loss function are then scaled by the number of gene sets S (denoted as *N*_S_) versus that of U (denoted as *N*_U_), and the number of spatial transcriptomic cells (denoted as *M*_T_) versus that of scRNA-seq cells (denoted as *M*_R_). The final loss is the sum of these two scaled losses:

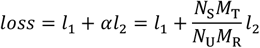

After feedforwarding each training batch twice, stPlus performs backward propagation with the final loss. stPlus optimizes parameters in the auto-encoder until convergence, and finally obtains cell embeddings of spatial transcriptomic data and reference scRNA-seq data. We also introduce two variations of stPlus, i.e. stPlus without the first part of loss (stPlus w/o P1) and stPlus without the second part of loss (stPlus w/o P2), to demonstrate how the two parts of loss affect the enhancement results.

In the predicting step, stPlus predicts the expression of spatially unmeasured genes using a strategy similar to SpaGE. For each spatial transcriptomic cell T_*i*_, stPlus calculates its cosine distance with each scRNA-seq cell R_*j*_ based on the learned cell embeddings. The neighboring 50 scRNA-seq cells are then used to predict the expression of unmeasured genes in cell T_*i*_ via a weighted *k*-NN approach. Specifically, stPlus filters out the neighbors with negative cosine similarity, and calculates the weight between T_*i*_ and *k*-th neighbor in the remaining *K* neighbors by:

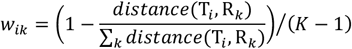

Finally, stPlus predicts expression of spatially unmeasured genes in cell T_*i*_ by the weighted expression of these genes in the *K* neighbors from reference scRNA-seq data:

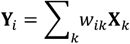

where **X**_*k*_ denotes expression data of genes to predict in the *k*-th neighbor. Leveraging the ensemble learning strategy, stPlus automatically adopts five epochs with minimal loss when training the auto-encoder, and averages the prediction results to achieve better and stable performance.

### 2.2 Data collection and preprocessing

We collected the five benchmarking dataset pairs adopted in SpaGE (Abdelaal, et al., 2020) to evaluate the enhancement performance of different computational methods. As shown in Table 1, the dataset pairs have diverse gene detection sensitivity levels, sample sizes, and number of spatially measured genes. Specifically, the five dataset pairs are made up of three spatial transcriptomic datasets (osmFISH, MERFISH and STARmap) from different mouse brain regions, and four reference scRNA-seq datasets (Zeisel, AllenVISp, AllenSSp and Moffit). The osmFISH dataset was retrieved from http://linnarssonlab.org/osmFISH/ (Codeluppi, et al., 2018). The MERFISH dataset was downloaded from https://doi.org/10.5061/dryad.8t8s248 (Moffitt, et al., 2018). The STARmap dataset is available at https://www.starmapresources.com/data (Wang, et al., 2018). Note that the labels of cell populations in the osmFISH and MERFISH datasets are accessible, while that of the STARmap dataset cannot be successfully aligned with cells. For the reference scRNA-seq data, the Zeisel dataset, which is provided by the same lab as that of the osmFISH dataset, was downloaded from http://linnars-sonlab.org/cortex/ (Zeisel, et al., 2015). The AllenVISp (Tasic, et al., 2018) and AllenSSp (Chatterjee, et al., 2018) datasets collected from https://portal.brain-map.org/atlases-and-data/rnaseq were more deeply sequenced than the Zeisel dataset. The AllenVISp was measured from a different brain region than the osmFISH dataset, while the AllenSSp dataset was measured from the somatosensory cortex, similar to the osmFISH dataset. The Moffit dataset, which was published in the same study of MERFISH, was retrieved from NCBI Gene Expression Omnibus (GEO) with the accession number GSE113576 (Moffitt, et al., 2018).

**Table 1.** Summary of the five benchmarking dataset pairs.

| Spatial/scRNA-seq dataset pair | Spatial transcriptomic data |  |  |  | Reference scRNA-seq data |  |  |  |
| --- | --- | --- | --- | --- | --- | --- | --- | --- |
|  | # of cells | # of genes | Data sparsity | Tissue | # of cells | # of genes | Data sparsity | Tissue |
| osmFISH_Zeisel | 3,405 | 33 | 29.7% | SMSc | 1,691 | 15,075 | 78.9% | SMSc |
| osmFISH_AllenVISp | 3,405 | 33 | 29.7% | SMSc | 14,249 | 34,617 | 74.7% | VISc |
| osmFISH_AllenSSp | 3,405 | 33 | 29.7% | SMSc | 5,577 | 30,527 | 69.8% | SMSc |
| MERFISH_Moffit | 64,373 | 155 | 60.6% | POR | 31,299 | 18,646 | 85.6% | POR |
| STARmap_AllenVISp | 1,549 | 1,020 | 79.0% | VISc | 14,249 | 34,617 | 74.7% | VISc |

Note: SMSc denotes somatosensory cortex; VISc denotes visual cortex; POR denotes pre-optic region.

For the implementation of stPlus, we followed the data preprocessing procedure of SpaGE except for the final *z*-score step (Abdelaal, et al., 2020). To be more specific, we filtered out the blank genes and the *Fos* gene (non-numerical values), and filtered out the cells labeled as ‘Ambiguous' in the MERFISH dataset. We only kept the cells from cortical regions in the osmFISH dataset. Each spatial transcriptomic dataset was then normalized via dividing the counts within each cell by the total number of transcripts within that cell, scaled by the median number of transcripts per cell, and log-transformed with a pseudo-count. For the reference scRNA-seq datasets, we filtered out genes expressed in less than ten cells. We also filtered out low-quality cells in the AllenVISp dataset according to the metadata (‘Low Quality’ and ‘No Class' cells), and filtered out the hippocampus cells in the Zeisel dataset according to the metadata. The filtered scRNA-seq datasets were next independently normalized by dividing the counts within each cell by the total number of transcripts within that cell, scaled by 10e6, and log-transformed with a pseudo-count.

### 2.3 Evaluation approaches

Considering that the technical challenge for imaging-based profiling of spatially resolved transcriptomics is to balance the number of genes imaged and the gene detection sensitivity, computational methods for data enhancement are anticipated to 1) accurately predict expression of unmeasured genes, especially for the data with small number of genes imaged, and 2) effectively impute expression of imaged genes to better identify cell populations, especially for the data with low gene detection sensitivity. We hence conduct a comprehensive evaluation of enhancement performance from two perspectives.

Firstly, we evaluated the prediction performance of gene expression by gene-wise and cell-wise Spearman correlation coefficients between the measured and predicted spatial profiles. The Spearman correlation coefficient over cells for each gene (gene-wise) can directly reflect the correlation between predicted and measured spatial profiles since the prediction object is a gene. The gene-wise coefficient was adopted as the primary evaluation metric in the two state-of-the-art methods specifically designed to enhance the spatial transcriptomic data (Abdelaal, et al., 2020; Lopez, et al., 2019). Given that cell heterogeneity is essential to single-cell studies, we additionally assessed cellular characteristics maintained in the predicted spatial profiles by Spearman correlation coefficients over genes for each cell (cell-wise) based on the idea that higher correlation per cell can better maintain cellular characteristics. We obtained the predicted spatial transcriptomics of all genes iteratively in cross-validation experiments, and then calculated Spearman correlation coefficients at cell-wise.

Secondly, we evaluated performance for the identification of cell populations by four clustering metrics. It is intuitive that, based on the ground-truth cell labels and the enhanced spatial transcriptomic data, a higher score of cell clustering metric indicates better performance for the identification of cell populations, and thus a better enhancement of spatial transcriptomics. Therefore, we assess clustering results based on the computationally predicted spatial transcriptomics, using clustering results based on the spatially profiled data as the baseline. Specifically, we used the standard pipeline with default parameters in Scanpy (Wolf, et al., 2018), a widely-used Python library for the analysis of single-cell data, to perform dimension reduction and cell clustering. We adopted a recently suggested clustering strategy for benchmark studies (Chen, et al., 2019). The strategy is based on Louvain clustering, a community detection-based clustering method (Blondel, et al., 2008; Levine, et al., 2015; Wolf, et al., 2018), and uses a binary search to tune the resolution parameter in Louvain clustering to make the number of clusters and the number of ground-truth cell labels as close as possible.

### 2.4 Metrics for assessment of clustering results

The clustering results were evaluated based on four widely-used metrics: adjusted mutual information (AMI), adjusted Rand index (ARI), homogeneity (Homo), and normalized mutual information (NMI). A comparison of ARI, NMI, and AMI was presented in (Romano, et al., 2016). Rand index (RI) represents the probability that the predicted clusters and the true cell labels will agree on a randomly chosen pair of cells. ARI is an adjusted version of RI, where it adjusts for the expected agreement by chance. ARI is preferred when there are large equal-sized clusters (Romano, et al., 2016). Both NMI and AMI are based on mutual information (MI), which assesses the similarity between the predicted clusters and true cell labels. NMI scales MI to be between 0 and 1, while AMI adjusts MI by considering the expected value under random clustering. AMI is theoretically preferred to NMI, even though NMI is also very widely-used and usually provides roughly the same results. Compared with ARI, AMI is preferred when the sizes of clusters are unbalanced and when there are small clusters (Romano, et al., 2016). We note that the sizes of cell populations in most spatial transcriptomic data, including the datasets used in this study, are unbalanced since some rare cell populations exist. Therefore, AMI is more appropriate in most cases, while ARI should be adopted when the clusters have nearly equal-sizes. The homogeneity score assesses whether the obtained clusters contain only cells of the same population, and it equals 1 if all the cells within the same cluster correspond to the same population.

To be more specific, suppose *T* is the known ground-truth labels of cells, *P* is the predicted clustering assignments, *N* is the total number of single cells, *x_i_* is the number of cells assigned to the *i*-th unique cluster of *P*, *y_j_* is the number of cells that belong to the *j*-th unique label of *T*, and *n_ij_* is the number of overlapping cells between the *i*-th cluster and the *j*-th unique label. Then the ARI score is computed as follows:

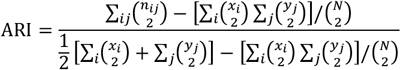

The NMI score is computed as the following formula:

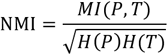

where *H*(·) is the entropy function, and *MI*(·,·) is the mutual entropy.

The AMI score is calculated as follows:

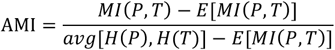

where *E*(·) denotes the expectation function.

The homogeneity score is computed as follows:

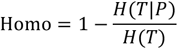

where *H*(*T*|*P*) denotes the uncertainty of true labels based on the knowledge of clustering assignments.

### 2.5 Baseline methods

We compared the performance of stPlus with four baseline methods, including SpaGE (Abdelaal, et al., 2020), Seurat (Stuart, et al., 2019), Liger (Welch, et al., 2019), and gimVI (Lopez, et al., 2019), with their default parameters or settings provided in the accompanying examples. Source code for implementing the baseline methods was obtained from the study of SpaGE, which is the first to systematically benchmark existing methods using the five dataset pairs introduced above (Abdelaal, et al., 2020). Data processing procedures, such as normalization and scaling, were also performed following the source code of each method.

## 3 Results

### 3.1 stPlus accurately predicts spatial transcriptomic data

To demonstrate the advantage of stPlus for predicting spatial gene expression, we conducted a series of 5-fold cross-validation experiments using five dataset pairs of diverse gene detection sensitivity levels, sample sizes, and number of spatially measured genes (Table 1). On each dataset pair, we randomly split genes shared between the two datasets into 5 folds, and predicted expression of the genes in each fold by the stPlus model trained with the genes in the remaining four folds. The predicted spatial transcriptomics of all genes in all cells can be obtained iteratively in the cross-validation experiments.

We first compared the Spearman correlation coefficients at gene-wise of stPlus to that of other methods. The gene-wise coefficients can directly reflect the correlation between predicted and measured spatial profiles since the prediction object is a gene. For each comparison, we conducted one-sided paired Wilcoxon signed-rank tests to test if stPlus achieves significantly higher Spearman correlation coefficients than the baseline methods at gene-wise. As shown in Fig. 2, stPlus consistently and significantly outperformed Seurat and Liger across all datasets. On the osmFISH_Zeisel dataset pair, stPlus significantly outperformed SpaGE (one-sided paired Wilcoxon test *p*-value < 0.01), and greatly improved the median Spearman correlation coefficient by 61.0% than gimVI although the *p*-value is not less than 0.01 (Fig. 2a). On the osmFISH_AllenVISp dataset pair, stPlus improved the median Spearman correlation coefficient by 11.2% than SpaGE, while slightly outperformed gimVI (Fig. 2b). On the osmFISH_AllenSSp dataset pair, stPlus again significantly outperformed SpaGE, and improved the median Spearman correlation coefficient by 7.6% than gimVI (Fig. 2c). On the above three dataset pairs, Seurat provided overall the lowest performance with Spearman correlation coefficients close to 0, which is consistent with the observations in the study of SpaGE (Abdelaal, et al., 2020), and suggests that the performance of Seurat heavily decreases when there are very few shared genes. On the MERFISH_Moffit dataset pair, which contains over 95 thousand cells, stPlus consistently achieved significantly better performance than other methods with at least 8.8% improvement of the median Spearman correlation coefficient (Fig. 2d). On the STARmap_AllenVISp dataset pair, stPlus significantly outperformed gimVI, while SpaGE achieved comparable performance (Fig. 2e). This is consistent with the observation in the study of SpaGE (Abdelaal, et al., 2020) that gimVI performs worse on data with low gene detection sensitivity. We note that there might exist inconsistencies between the results of 5-fold cross-validation in this study and the results of leave-one-out cross-validation in the study of SpaGE (Abdelaal, et al., 2020), which simplifies the task because the training set contains more genes highly correlated with the gene to predict.

**Fig. 2.**
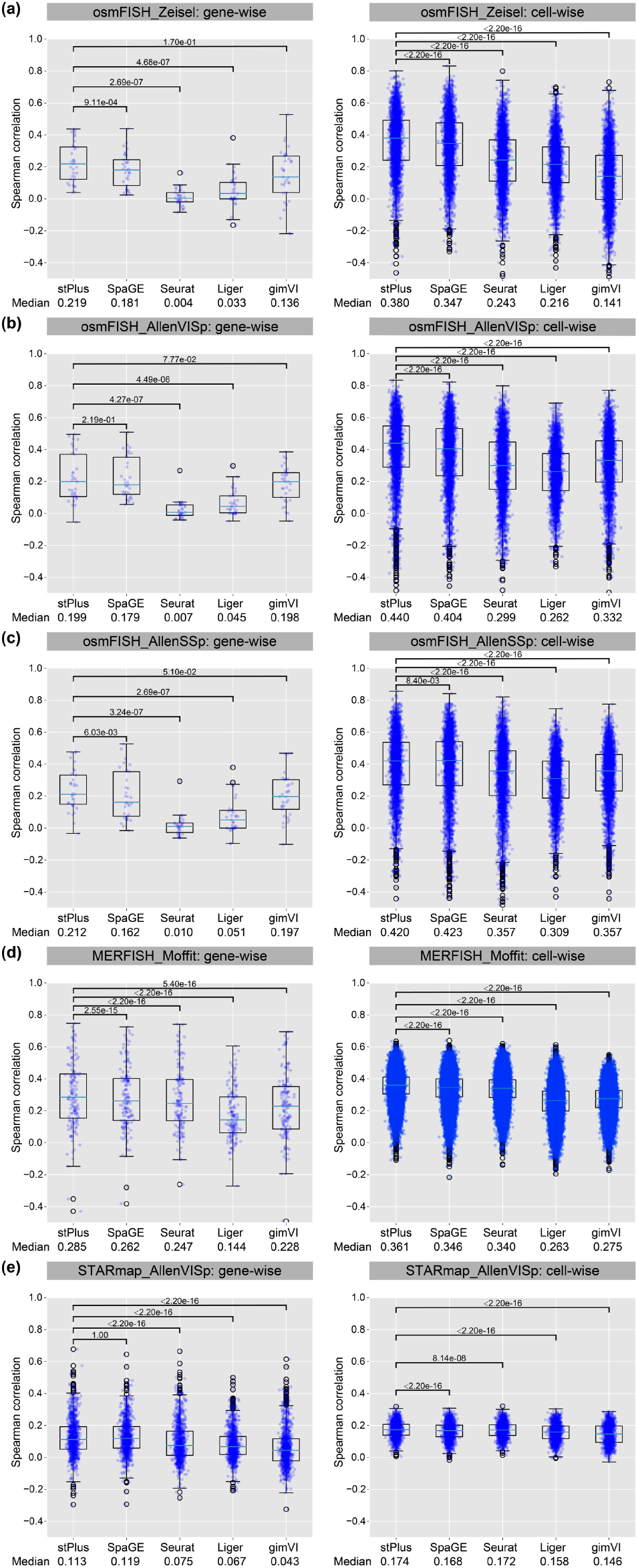
Spearman correlation coefficients of different methods on various dataset pairs. The median Spearman correlation coefficient and *p*-value of one-sided paired Wilcoxon signed-rank tests are reported.

We then compared the Spearman correlation coefficients of different methods at cell-wise. The cell-wise coefficients can reflect cellular characteristics maintained in the predicted spatial profiles. For each comparison, we also conducted one-sided paired Wilcoxon signed-rank tests to test if stPlus achieves significantly higher Spearman correlation coefficients at cell-wise. As shown in Fig. 2, stPlus consistently achieved significantly higher coefficients than all the four baseline methods across all the five dataset pairs (one-sided paired Wilcoxon test *p*-value < 0.01), and has an average improvement of 23.2% on the median Spearman correlation coefficient over the second-best method. All these results suggest the superior performance of stPlus for the prediction of spatially unmeasured gene expression. Compared with stPlus w/o P1, stPlus improved the median gene-wise and cell-wise Spearman correlation coefficients by an average of 2.9% and 0.1%, respectively. However, compared with stPlus w/o P2, the average improvements are 8.9% and 7.5%, respectively, which suggests that the accurate prediction of gene expression mainly benefits from the second part of loss function.

### 3.2 stPlus facilitates the identification of cell populations

The essential step in single-cell analyses is the characterization of known or novel cell types, which is based on the accurate identification of cell populations. However, in technical limitations, either small number of genes imaged or low gene detection sensitivity constitutes major hindrances to the identification of cell populations in spatial transcriptomic data. Therefore, we used the measured spatial transcriptomic data as the baseline to verify if the computationally enhanced spatial transcriptomics can better identify cell populations. Specifically, we obtained cell labels of the osmFISH and MERFISH datasets from the original studies, while that of the STARmap dataset cannot be successfully aligned with cells. We then used the cell labels as ground truth, and evaluated the performance by four clustering metrics, including AMI, ARI, Homo, and NMI. It is intuitive that, based on the enhanced data, a higher score of clustering metric indicates better performance for the identification of cell populations, and thus a better enhancement of spatial transcriptomics. Note that the sizes of different cell populations in most spatial transcriptomic data, including the datasets used in this study, are unbalanced since there exist some rare cell populations, and AMI is preferred in this case compared with ARI (Romano, et al., 2016).

Firstly, we performed cell clustering using the predicted gene expression in each fold of the 5-fold cross-validation, which means that only one-fifth of the genes are used for clustering. As shown in Fig. 3a, clustering performance using the originally profiled spatial transcriptomic data is unsatisfactory and fluctuates greatly, which is expected since few genes were used (e.g., about six genes for the osmFISH dataset). However, the clustering performance can be significantly improved using the computationally predicted gene expression. For example, SpaGE and gimVI achieved better clustering performance with smaller variance on the first three dataset pairs that are based on the osmFISH dataset. Consistent with the performance evaluated by Spearman correlation coefficients, Seurat provided the lowest clustering performance on the first three dataset pairs, while the advantages of Seurat over other methods were observed on the MERFISH dataset, which again suggests that the performance of Seurat is greatly affected by the number of genes shared between two datasets. Among these four dataset pairs, stPlus achieved the overall best clustering performance, especially on the first three dataset pairs (Fig. 3a).

**Fig. 3.**
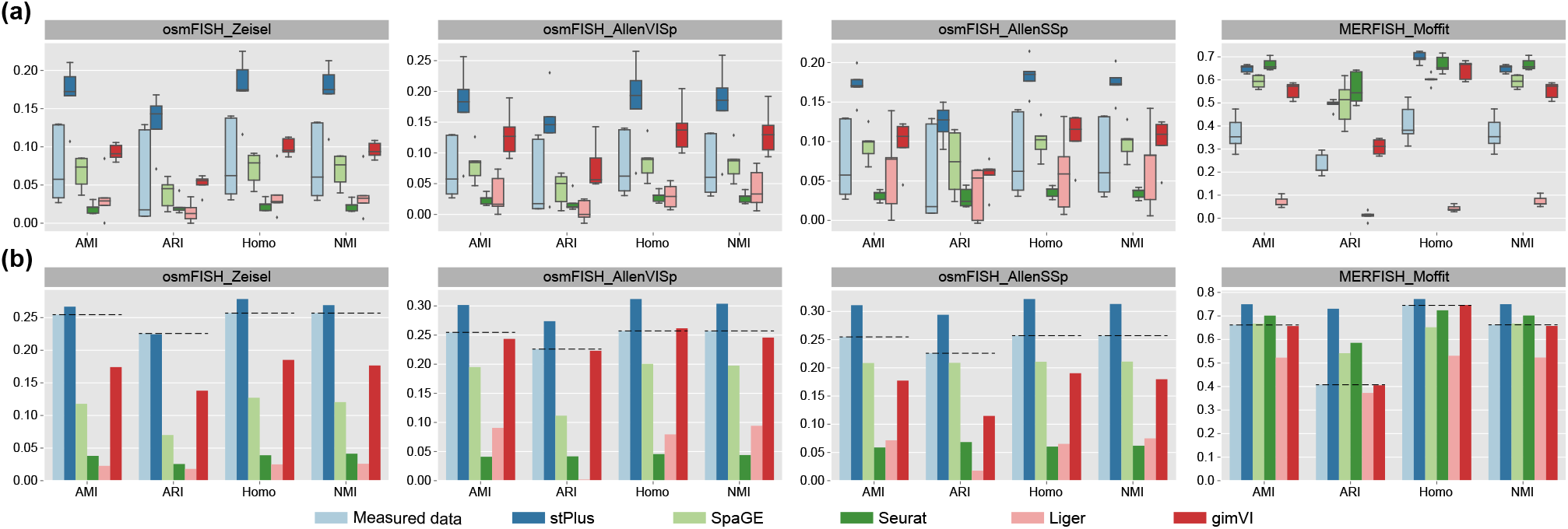
Clustering results of spatial transcriptomic cells evaluated by AMI, ARI, Homo, and NMI. (a) Clustering results using the measured or computationally predicted expression of genes in each fold of the 5-fold cross-validation. (b) Clustering results using the measured or computationally predicted expression of all shared genes.

Secondly, we obtained the predicted spatial transcriptomics of all genes iteratively in the cross-validation experiments. The conduction of cross-validation experiments can be regarded as a strategy for data enhancement. As expected, the baseline performance (dotted lines in Fig. 3b) was sig-nificantly improved using the data with all genes. On the first three dataset pairs, only when using the gene expression data predicted by stPlus can the clustering performance exceed the baseline. On the MERFISH_Moffit dataset pair, stPlus again outperformed the baseline and other computational methods, while Seurat also achieved better clustering performance than the baseline. These results not only suggest that stPlus is capable of predicting spatially unmeasured gene expression, but also demonstrate that the data enhanced by stPlus can provide superior performance for the identification of cell populations than that enhanced by existing methods and even the originally profiled spatial transcriptomic data. Compared with stPlus w/o P1, stPlus improved AMI, ARI, Homo, and NMI by an average of 7.9%, 21.0%, 6.3%, and 7.8%, respectively. However, compared with stPlus w/o P2, the average improvements are 5.6%, 3.4%, 4.0%, and 5.5%, respectively, which suggests that the effective characterization of cell heterogeneity mainly benefits from the first part of loss function.

### 3.3 stPlus is scalable to large datasets

Recent efforts of cell atlas consortiums have generated massive amounts of scRNA-seq data, providing a wealth of reference data on the diversity of cell types across organisms, developmental stages, and disease states (Davie, et al., 2018; Han, et al., 2018; Rozenblatt-Rosen, et al., 2017; Tabula Muris, et al., 2018; Zeisel, et al., 2018; Zheng, et al., 2017). Besides, high-throughput technologies now allow the simultaneous profiling of massive cells. Therefore, computational efficiency and scalability are important for a computational method to facilitate the enhancement of spatial transcriptomics. We have demonstrated the superior enhancement performance of stPlus on the MERFISH_Moffit dataset pair, which is composed of 64,373 spatial transcriptomic cells and 31,299 scRNA-seq cells. To assess the computational efficiency and scalability of different methods, we benchmarked the average running time and the peak memory usage in experiments of the 5-fold cross-validation on the MERFISH_Moffit dataset pair. The experiments were run on a machine with an Intel Xeon E5-2660 v4 X CPU, a GeForce GTX 1080 and 500GB of RAM on CentOS 7 operating system. As shown in Table 2, stPlus provided satisfactory computational efficiency and scalability in large data, while SpaGE achieved the best computational efficiency. The efficiency advantage of SpaGE mainly benefited from the joint embedding step, where it is based on principal component analysis while stPlus needs iterative training. Seurat, Liger, and gimVI performed relatively worse for the computational efficiency. Note that gimVI, a probabilistic method, directly performs prediction without the embedding step. The peak memory usage of stPlus is moderate among all the methods, and is reasonable since stPlus additionally incorporated genes unique to the reference 31,299 scRNA-seq cells.

**Table 2.**
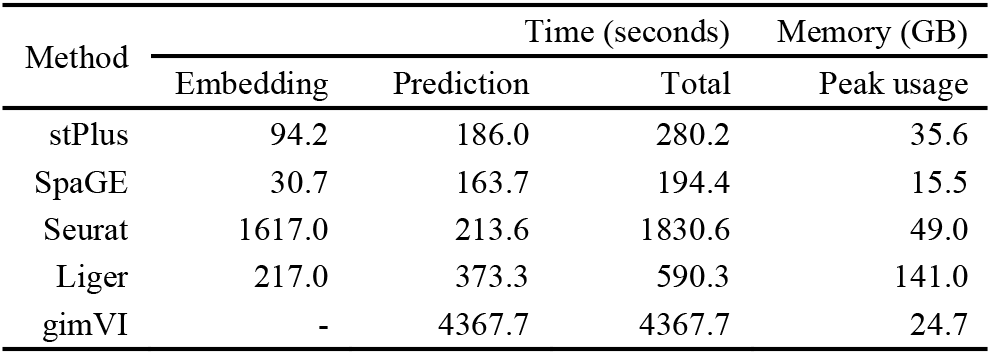
Average running time and peak memory usage in experiments of the 5-fold cross-validation on the MERFISH_Moffit dataset pair.

Data visualization is fundamental to single-cell analyses by providing an intuitive understanding of cellular composition. To test if the predicted spatial transcriptomics of large data can provide satisfactory data visualization results, we used the standard pipeline with default parameters in Scanpy (Wolf, et al., 2018) to perform dimension reduction and UMAP visualization (McInnes, et al., 2018). As shown in Fig. 4, compared to other methods, using the expression of spatial genes enhanced by stPlus, the patterns of various cell types can be better re-characterized, and some biological variations are even better distinguished than using the measured data. For example, the excitatory and inhibitory cells can be better separated using the expression data enhanced by stPlus. To assess the identification of minor cell types quantitatively, we left out the cell types that account for more than 10% (Inhibitory: 38.5%, Excitatory: 18.3%, and Astrocyte: 13.0%), resulting in more balanced and minor cell types. We evaluated the clustering performance of different methods. stPlus again provided better performance than the measured data. Compared with the second-best method, stPlus improved AMI, ARI, Homo, and NMI by 3.8%, 13.3%, 7.4%, and 3.8%, respectively. All the above observations suggest that stPlus can not only offer satisfactory computational efficiency and memory usage, but also provide the scalability to visualization of large data based on the enhanced spatial transcriptomics.

**Fig. 4.**
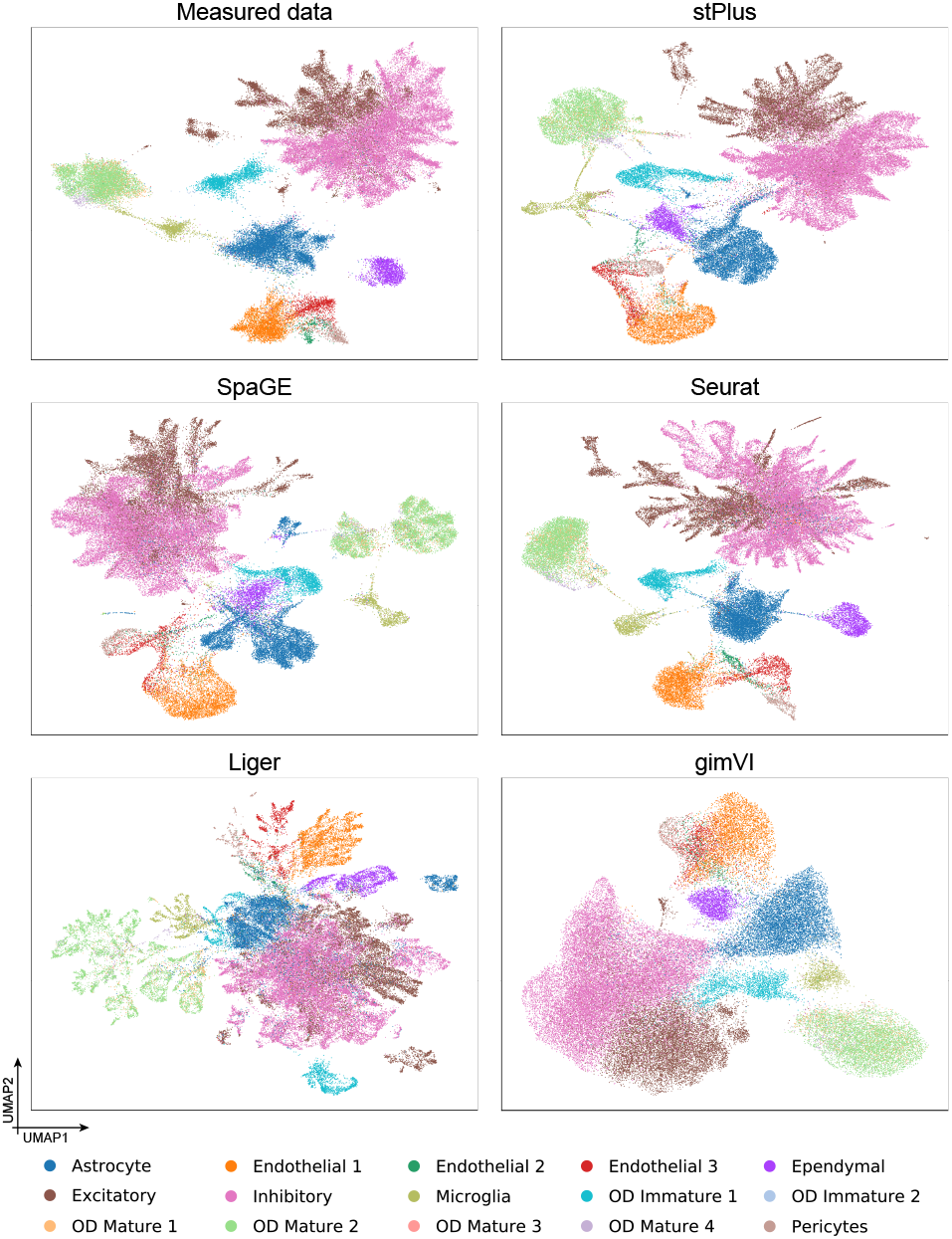
UMAP visualization of cells in the MERFISH dataset using the measured or computationally predicted expression of all shared genes.

### 3.4 stPlus is robust to the choice of hyperparameters

The major hyperparameters of stPlus are the number of neighboring cells used to predict, and the number of epochs adopted to perform ensemble learning. In this section, we further used the MERFISH_Moffit dataset pair to demonstrate the robustness of stPlus to the choice of these two hyperparameters. We note that in the MERFISH dataset, the dominant cell types, i.e., excitatory and inhibitory cells, constitute over 56.7% of the total cells, while the rare cell types, i.e., OD Immature 2, OD Mature 3, and OD Mature 4 cells, constitute less than 0.8%. Therefore, compared with ARI, AMI is preferred for the assessment of cell clustering on this dataset (Romano, et al., 2016). When we varied one hyperparameter, we fixed the other parameters to the default setting. We used the spatial transcriptomics of all genes iteratively predicted in the 5-fold cross-validation experiments to evaluate the robustness of stPlus. As shown in Fig. 5a, stPlus achieved stable gene-wise and cell-wise Spearman correlation coefficients with different choices of the number of neighboring cells used to predict. Besides, there are no significant fluctuations in the clustering metrics except ARI. We next varied the number of epochs adopted to perform ensemble learning. As shown in Fig. 5b, the performance assessed by gene-wise and cell-wise Spearman correlation coefficients is stable, and the clustering metrics show no significant fluctuations except ARI. The results indicate that stPlus is robust to the choice of hyperparameters.

**Fig. 5.**
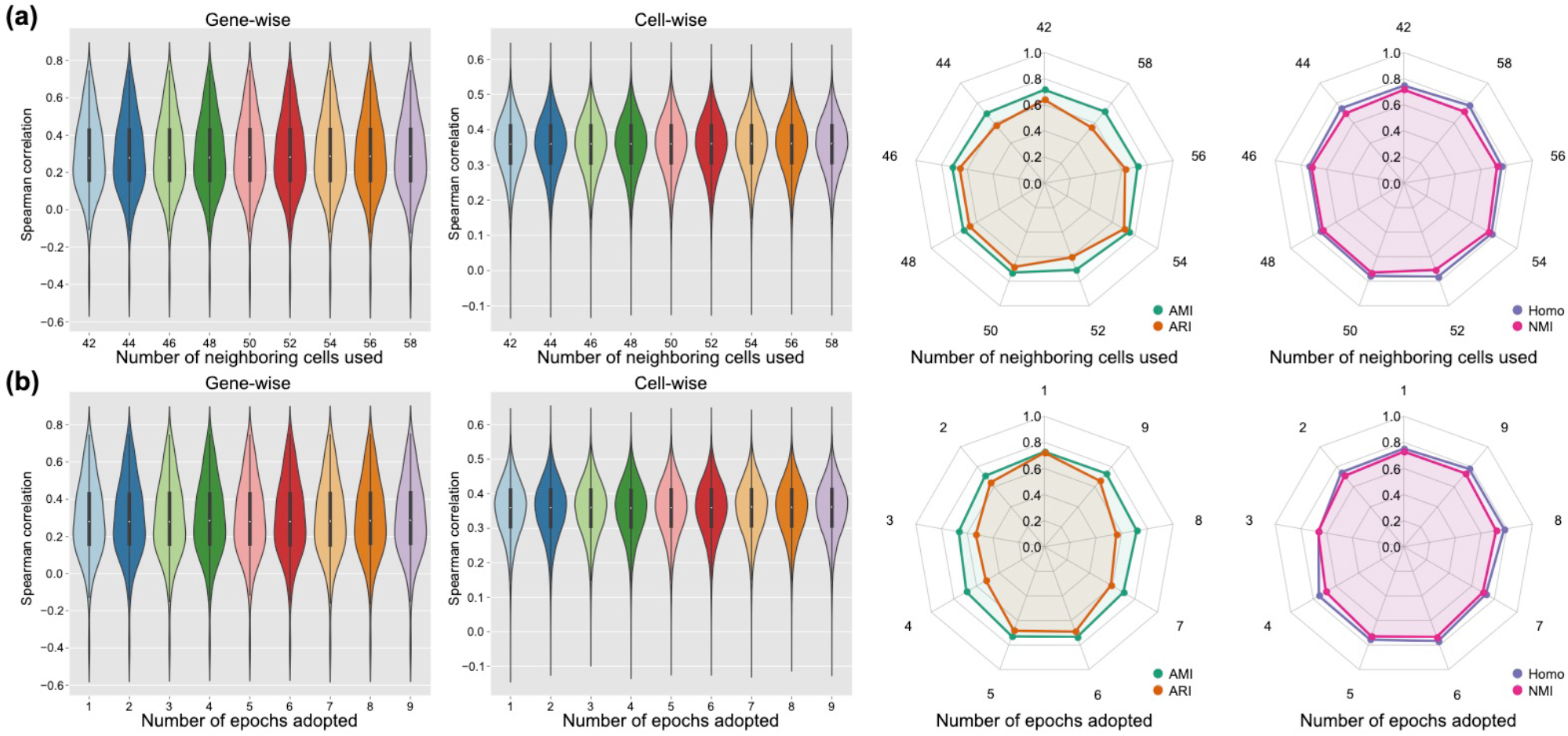
The robustness of stPlus to the choice of (a) number of neighboring cells used to predict, and (b) number of epochs adopted to perform ensemble learning.

### 3.5 Genes unique to reference data contain cell heterogeneity

The above assessments are based entirely on the profiled spatially resolved transcriptomics. To verify if the genes inherent to reference scRNA-seq data can characterize the cellular composition, instead of 5-fold cross-validation, we further used all shared genes to train models and predicted expression of the genes unique to scRNA-seq data. Since expression of these genes was not measured in spatial transcriptomic cells, we only evaluated the clustering performance using the predicted expression of these genes, and compared it with the clustering performance using the predicted expression of genes shared between spatial and scRNA-seq datasets, i.e., the results in Fig. 3b. The measured spatial transcriptomic data was again used as the baseline (light blue markers and dotted lines in Fig. 6). As shown in Fig. 6, most methods provided better clustering performance on the first three dataset pairs using the genes unique to scRNA-seq data than using the 33 shared genes. However, on the MERFISH_Moffit dataset pair that has more shared genes, most methods achieved better clustering performance using the 153 shared genes, while stPlus offered comparable performance using the genes unique to scRNA-seq data. In addition, stPlus outperformed other methods by all the four metrics across all the four dataset pairs, which indicates that the spatial expression of genes unique to scRNA-seq data predicted by stPlus will continue to deepen our understanding of the characterization of cell heterogeneity.

**Fig. 6.**
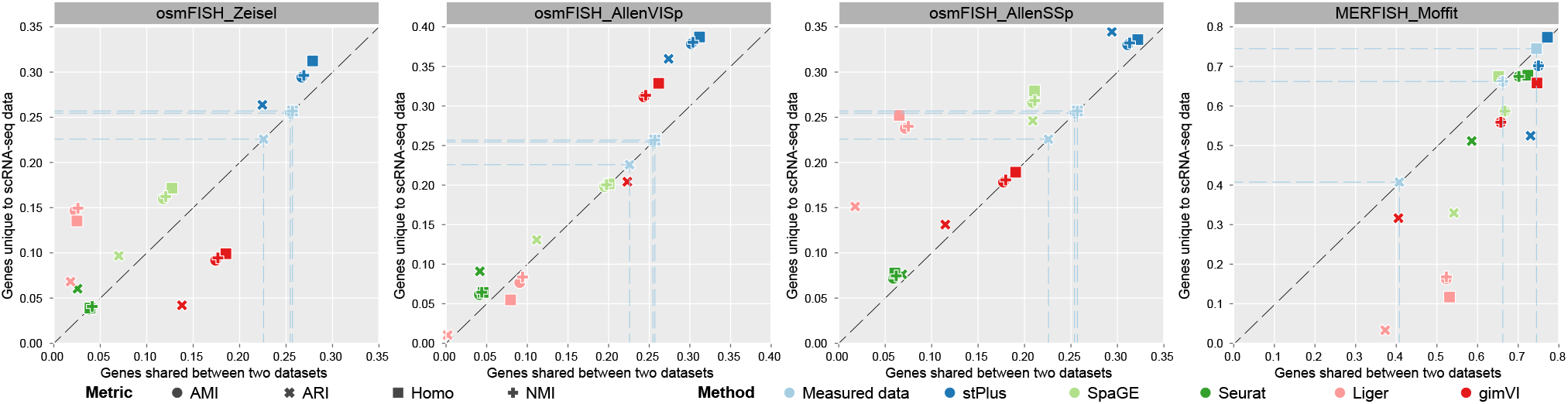
Clustering results of spatial transcriptomic cells using the predicted expression of genes shared between two datasets versus genes unique to scRNA-seq data.

We further demonstrated that the predicted expression of scRNA-seq-unique genes can provide biological insight into the identified cell population. Taking the MERFISH_Moffit dataset pair as an example, we used Wilcoxon rank-sum test in Scanpy (Wolf, et al., 2018) to find the top 5 differentially expressed genes (DEGs) for each cluster. We performed Gene Ontology (GO) enrichment analysis using the DEGs uniquely obtained by each method (Ashburner, et al., 2000; Gene Ontology, 2021). Using the DEGs uniquely obtained by stPlus, the significantly enriched results (generation of neurons, axon development, nervous system development, cell development, neuron differentiation, and neurogenesis) are related to the biological process of neurons, which is consistent with the original spatial data. However, none of the significantly enriched biological process using the DEGs uniquely obtained by each of other methods is directly related to the original spatial data. These results indicate that stPlus opens a new avenue for the identification of marker genes and further provides biological insight into spatial transcriptomics analysis.

## 4 Discussion

In this work, we proposed stPlus for the accurate enhancement of spatial transcriptomics, and introduced a clustering-based approach for the systematical assessment of enhancement performance. stPlus simultaneously models the genes shared between spatial transcriptomic data and reference scRNA-seq data and the genes unique to scRNA-seq data. We demonstrated that stPlus outperforms baseline methods in accurately predicting expression of unmeasured genes. Besides, stPlus facilitates the identification of cell populations by enhancing spatial transcriptomics. The predicted spatial expression of scRNA-seq-unique genes also provides potential for the characterization of cell heterogeneity. In addition, stPlus is robust and scalable to dataset pairs of diverse gene detection sensitivity levels, sample sizes, and number of spatially measured genes. We also provided user-friendly interfaces, detailed documents, and quick-start tutorials to facilitate the application of stPlus.

Certainly, our modeling framework is flexible and can be extended easily. Firstly, gene expression data used in this study can be further combined with spatial coordinates. Secondly, we can incorporate other types of profiles, e.g., epigenetic profiles, as the additional reference data (Chen, et al., 2020; Chen, et al., 2020). Thirdly, we can also extend stPlus by incorporating advanced deep neural networks to capture higher-level features of the profiled cells in spatial transcriptomic data (Liu, et al., 2020).

## Acknowledgements

We appreciate the authors of SpaGE for the benchmarking data and code.

